# Resource partitioning in organosulfonate utilization by free-living heterotrophic bacteria during a North Sea microalgal bloom

**DOI:** 10.1101/2024.12.10.627767

**Authors:** Chandni Sidhu, Daniel Bartosik, Vaikhari Kale, Anke Trautwein-Schult, Dörte Becher, Thomas Schweder, Rudolf I. Amann, Hanno Teeling

## Abstract

Blooming microalgae (phytoplankton) release diverse organic molecules that fuel the marine pools of dissolved and particulate organic matter. A highly specialized community of heterotrophic bacteria rapidly remineralizes substantial parts of this organic matter in the sun-lit upper ocean. In particular, microalgae produce large quantities of various organosulfur compounds that can serve as carbon and sulfur sources for bacteria.

Here, we report on the analyses of a time series of previously generated 30 long-read metagenomes, 30 corresponding deeply sequenced short-read metatranscriptomes and 15 metaproteomes from 0.2-3 µm size fractions that we sampled in 2020 during a biphasic phytoplankton bloom in the German Bight (Southern North Sea). We analyzed the assembled contigs as well as 70 bacterial metagenome-assembled genomes that recruited the highest transcript numbers with respect to the utilization of methyl sulfur compounds (dimethylsulfoniopropionate (DMSP), dimethyl sulfide (DMS), dimethyl sulfone (DMSO_2_)), C3-sulfonates (2,3-dihydroxypropane-1-sulfonate (DHPS), 3-sulfolactate, 3-sulfopyruvate) and 2-aminoethanesulfonic acid (taurine).

We observed a pronounced resource partitioning among bacterial clades that utilize distinct organosulfur compounds, which may explain successions of these clades during the studied bloom. *Alphaproteobacteria* were the most active and degraded a variety of organosulfonates via various metabolic routes. However, we also found previously underreported roles of members of the *Bacteroidota* and *Gammaproteobacteria* as efficient degraders of DMSP, DMS, and DMSO_2_. One striking observation was a strong preference for DMSP cleavage in *Bacteroidota* as opposed to DMSP demethylation in *Alphaproteobacteria* and indications for a particular proficiency for taurine utilization in Ilumatobacter_A and *Acidimicrobiia*.

**Importance:** Sulfur-containing low-molecular-weight algal metabolites play an important role in overall marine carbon and sulfur fluxes. This study highlights that such compounds may play a crucial role in governing the succession of distinct bacterioplankton clades in response to phytoplankton blooms in coastal shelf areas of the temperate zone, such as the German Bight of the North Sea. While *Alphaproteobacteria* are the most versatile and competitive degraders of dissolved organosulfur compounds during such blooms, this study repositions clades previously thought to play only a more limited role in dissolved organosulfur metabolism *in situ*, such as *Gammaproteobacteria*, *Bacteroidota*, and *Acidimicrobiia*, as crucial contributors to the remineralization of organosulfur compounds in the upper ocean. This study also highlights the high level of interconnectedness of bacterial carbon and sulfur cycling during phytoplankton blooms.

## Introduction

Marine planktonic uni- to pluricellular microalgae (phytoplankton) account for about half of the global photosynthetic carbon dioxide fixation (1, 2). Their activities peak during sometimes vast bloom events that occur globally, at which in particular small cell-sized phytoplankters can reach high growth and carbon fixation rates (3). In temperate coastal areas, phytoplankton blooms often occur in spring, facilitated by accumulated inorganic nutrients from winter in combination with increasing water temperatures and solar irradiation. Blooming phytoplankton release diverse organic compounds that fuel the marine pools of dissolved and particulate organic matter (DOM, POM). A highly specialized community of mostly heterotrophic bacteria has adapted to and co-evolved with phytoplankton for millions of years (diatoms as youngest major phytoplankton lineage are at least 250 million years old (4)). While bacteria-phytoplankton interrelations are manifold and cover the full spectrum from beneficial (including symbiotic) functions (5–7) to antagonistic (competitive or even algicidal) functions (8), these bacteria all decompose algal-derived organic matter. Bacteria that thrive during phytoplankton blooms thereby exert a substantial influence over global carbon fluxes, as they determine the proportions of algal organic matter that are rapidly remineralized in surface waters versus those that are either sequestered in persistent DOM or sink out of the epipelagic zone as POM. The bulk of bloom-associated bacteria (often > 99%, e.g., (9)) are free-living (FL) and feed preferentially on DOM, but particle-attached (PA) bacteria carry out a gatekeeping function, as they are crucial for the mineralization and solubilization of a substantial proportion of the POM (10, 11).

Both, bloom-associated FL and PA microbial communities are dominated by *Alphaproteobacteria*, *Gammaproteobacteria* and *Flavobacteriia*^1^. However, different genera usually dominate both communities, often in swift successions that we observed to be more dynamic among PA bacteria in a recent study of an exemplary North Sea spring phytoplankton bloom (12). These bloom-associated bacteria form a tight-knit and in parts co-dependent food network that collectively remineralizes algal biomass, whereby distinct bacterial clades tend to specialize in distinct substrate classes. This resource partitioning mechanism represents not only an efficient division of labor but also serves as a means to minimize competition. Substrate chemical diversity and structural complexity can amplify resource partitioning, as is evident for the wealth of distinct polysaccharides that phytoplankters produce. The more complex a polysaccharide substrate, the more distinct genes are required for the involved complex uptake and breakdown protein machineries, which is why no bacterium can harbor the genes to decompose all algal polysaccharides. Thus, the decomposition of these polysaccharides is a collective endeavor of multiple polysaccharide specialists (12–14), at which members of the *Flavobacteriia* (15) and to a lesser extent *Gammaproteobacteria* (16) have been shown to excel.

Resource partitioning is not restricted to structurally complex, high molecular weight macromolecules, but is widespread and includes a plethora of structurally simple, low molecular weight substrates. This has been exemplarily demonstrated in an investigation of the metabolic responses of *Ruegeria pomeroyi* DSS-3 (*Alphaproteobacteria*), *Stenotrophomomas* sp. SKA14 (*Gammaproteobacteria*), *and Polaribacter dokdonensis* MED152 (*Flavobacteriia*) when co-cultured individually with the centric diatom *Thalassiosira pseudonana*. This study did uncover utilization of various amino acids/peptides, nucleosides/nucleotides, saccharides, vitamins, organic acids and dissolved organosulfur (DOS) compounds, of which only a few were likewise consumed by two or even all three bacterial model strains (17).

Among the abundant low molecular weight substrates that algae and other marine organisms produce are various DOS compounds such as dimethylsulfoniopropionate (DMSP), dimethyl sulfide (DMS), dimethyl sulfone (DMSO_2_) and 2-aminoethanesulfonic acid (taurine), and C3-sulfonates such as 3-sulfopropanediol (DHPS), 3-sulfolactate and 3-sulfopyruvate (18–21). Such DOS compounds are relevant in terms of both carbon and sulfur cycling, as they can for example make up a substantial proportion of the DOM released by zooplankton (21, 22) and even 20% to 40% of the total organic sulfur in marine sediments (23).

DMSP is particularly abundant, as it is not only produced by marine micro- and macroalgae but also by some higher plants (24), corals (25) and bacteria (18). It can account for up to 16% of the carbon in phytoplankters (26), and total global production ranges in the scale of one gigaton per annum (27–29). DMSP plays a vital role for many marine bacteria, not only as a source of carbon and energy but also of reduced sulfur. Members of the *Alphaproteobacteria* such as the SAR11/SAR116 (30, 31) and the *Roseobacter* clades (32, 33) are prolific DMSP degraders. DMSP utilization can occur either via demethylation or cleavage (34) (Fig. 1). The former involves the genes *dmdABCD* (along with a gene coding for the selenium-binding protein SELENBP1 (K17285)) and the key enzyme DmdA for the initial demethylation, which leads to methanethiol, and subsequently via sulfide and acetaldehyde to acetate. The latter involves DddP (35) or various alternative cupin superfamily DMSP lyase isoenzymes (DddK, DddL, DddQ, DddW, DddY and the recently discovered DddU (36–38)) and leads to DMS and acrylate. DMS is one of the major volatile organic sulfur compounds in the oceans and accounts for about 50% of the sulfur that enters the atmosphere via sea-to-air exchanges (39). DMS can be further oxidized to DSMO/DMSO_2_ and ultimately to methanesulfonate and sulfite (Fig. 1), further contributing to the formation of atmospheric cloud condensation nuclei (for review: (18)). The oxidation of DMS is facilitated by DmsBC (dimethyl sulfoxide reductase), and that of DSMO/DMSO_2_ by SsuE/SfnG (dimethylsulfone monooxygenase) and SsuD (alkanesulfonate monooxygenase).

**Figure 1.**
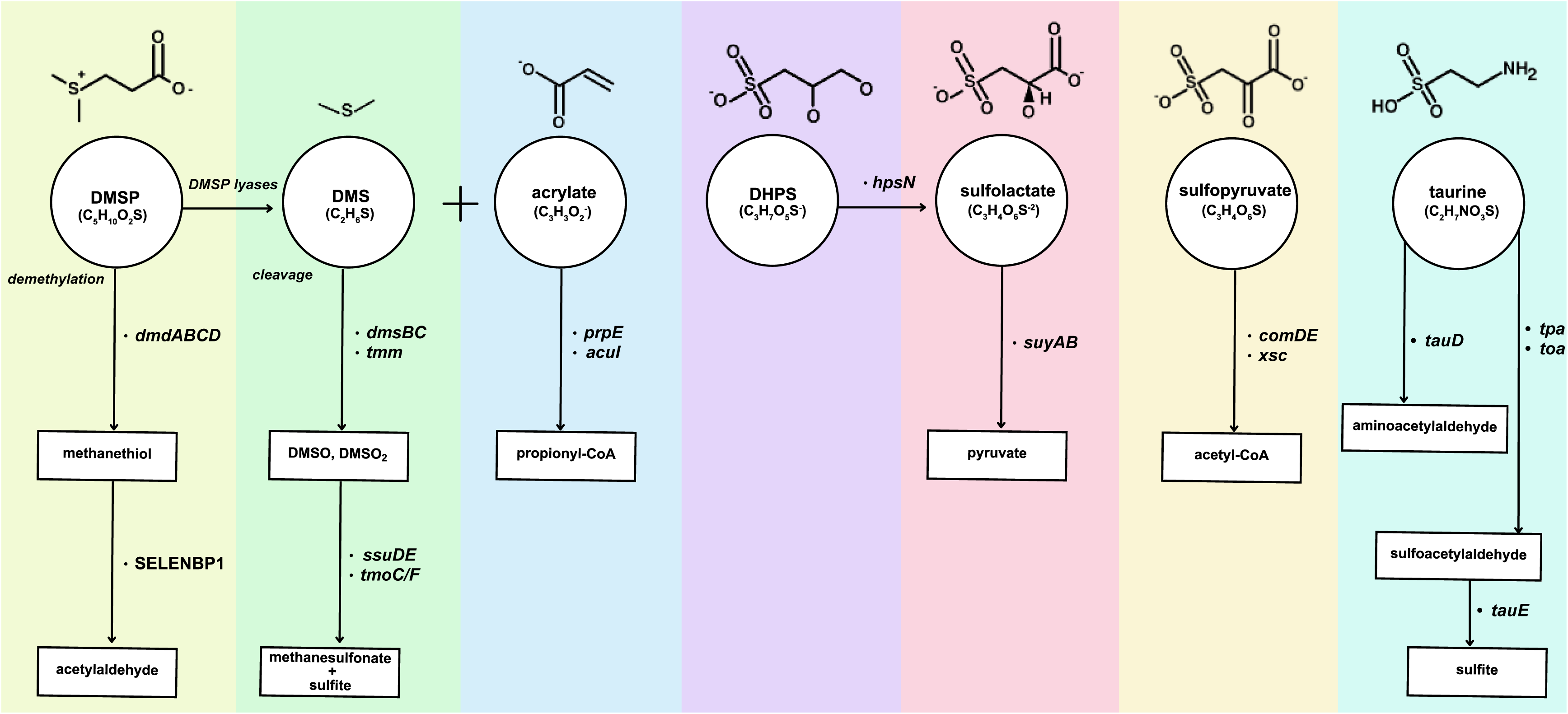
Organosulfonate compounds analyzed in this study. Identified genes in either assembled contigs or MAGs are indicated together with the conversions they catalyze. Key compounds include DMSP (dimethylsulfoniopropionate), DMS (dimethyl sulfide), DMSO (dimethyl sulfoxide), DMSO₂ (dimethyl sulfone), DHPS (2,3-dihydroxypropane-1-sulfonate), acrylate, sulfolactate, sulfopyruvate and taurine. More detailed descriptions of the enzymatic reactions are provided in the supplementary information (Table S2, S3).

Taurine, an amino acid-like compound, represents another abundant organosulfonate (40). It acts as an osmotic stress protectant in numerous marine metazoans including crustacean zooplankton and algae, where it regulates the internal osmotic balance and thereby shields against external salinity fluctuations (41). Besides, taurine may also partake in other physiological functions, including as detoxifying antioxidant and in macromolecular stabilization (42, 43). Thus, taurine plays multiple roles for the fitness of diverse marine organisms (44). When released, e.g., via zooplankton excretion, taurine contributes to the pool of DOS compounds and has been shown to be efficiently metabolized by dedicated bacterial clades such as the alphaproteobacterial SAR11 and, to a lesser extent, marine *Thaum-* and *Euryarchaeota* (45).

Besides DMSP and taurine, phytoplankters produce a number of other DOS compounds, including N-acetyltaurine, isethionate, choline-O-sulfate, cysteate, DHPS, and glutathione (e.g., (21, 32, 46)). However, owing to natural variability, our knowledge about the *in situ* relevance of these compounds as substrates for dedicated clades of marine heterotrophic bacteria and the associated resource partitioning strategies is still patchy.

In the year 2020, we sampled a spring phytoplankton bloom off Helgoland island in the German Bight (southern North Sea) at high temporal resolution. This has resulted in a comprehensive dataset of physicochemical data, microalgal microscopic counts and diversity data, as well as long-read metagenomes (n = 30) and corresponding short-read metatranscriptomes (n = 30), cell counts and CARD-FISH data of FL (0.2-3 µm size fraction) bacteria (13). Unlike our previous studies on spring blooms at this location (12–14, 16, 47, 48), the 2020 spring bloom featured two very distinct phases that were both almost entirely dominated by centric diatom species, the first by *Ditylum brightwellii*, and the second by *Chaetoceros* sp. and *Cerataulina pelagica* (for details, see (13)). The bacterioplankton response was characterized by a swift succession of various abundant clades, including *Aurantivirga* and *Polaribacter* (both *Flavobacteriia*), *Amylibacter* (*Alphaproteobacteria*) and *Glaciecola* (*Gammaproteobacteria*).

So far, we focused on the substrate niches of abundant, active, polysaccharide-degrading bacterioplankton clades (13). Here, we investigate a different angle of bacterial resource partitioning and explore FL clades of abundant, active non-polysaccharide specialists that expressed genes for the utilization of various methyl sulfur and other organosulfur compounds (Fig. 1). For this purpose, we complemented our dataset with a series of state-of-the-art metaproteomes (n = 15, protein groups = 21,472) that we previously analyzed in a different context (49, 50), in order to not only detect *in situ* gene transcription but where possible also actual protein abundance.

## Results

### Organosulfonate gene expression shifted between distinct bloom phases

Individual assemblies of all 30 metagenomes (1 PacBio Sequel IIe SMRT cell per sample) comprised a total of 80,322 contigs with a combined length of approximately 2.1 Gbp. Mapping of all 30 transcriptomes (100 M Illumina HiSeq 3000 PE reads per sample) onto all assemblies recruited 71.5% of the transcript reads, demonstrating the representativeness of the dataset.

Analysis of organosulfonate metabolism revealed genes coding for enzymes of various taurine utilization pathways, including (i) the oxidative decarboxylation to aminoacetaldehyde and sulfite by taurine dioxygenase (TauD, K03119) (51, 52), (ii) the oxidative transamination to sulfoacetaldehyde and ultimately acetylphosphate by a series of enzymes including taurine-pyruvate aminotransferase (Tpa, K03851), (iii) taurine-2-oxoglutarate transaminase (Toa, K15372), (iv) sulfoacetaldehyde acetyltransferase (Xsc, K03852) (53, 54), and (v) the dehydrogenation of taurine to sulfoacetaldehyde by TauXY (K07255, K07256) (55). Furthermore, we identified genes for the transformation of taurine to glutamyl-taurine by gamma-glutamyltranspeptidase (GGT, K00681 & K18592) (56), and for the conversion of taurine to taurocyamine by glycine amidinotransferase (GATM, K00613) (57).

We analyzed the expression of genes associated with these pathways over the course of entire studied bloom (Fig. 2A). On overall, *tauD* gene expression was higher during the pre-bloom and first bloom phases before April 30, decreased during the ensuing second bloom phase and ramped up again during the terminal phase (Fig. 2B). The expression of *ggt* followed a similar trend (Fig. 2B). Taxonomic analysis of *tauD* sequences revealed that *Alpha*- and *Gammaproteobacteria* were the major contributors to *tauD* expression in the pre- and first-bloom phases (Fig. 2C). In contrast, the rise in expression during the second and terminal bloom phases was mainly attributed to *Gammaproteobacteria*, while activities of *Alphaproteobacteria* decreased. We also analyzed the presence of the taurine-specific transporter genes *tauABC* and *tauT*. On overall, *tauABC* exhibited higher expression levels than *tauT* genes (Fig. 2D). Taxonomic sequence analyses revealed that the *tauABC* genes were mostly expressed in *Alphaproteobacteria*, whereas expression in *Gammaproteobacteria* was negligible (Fig. 2E). We also detected expression of the sulfite exporter *tauE*, which is used for the excretion of sulfite produced during the metabolism of C2-sulfonates (58). Expression of *tauE* was highest during the first bloom phase, similar to the above-mentioned taurine transporters *tauABC* and *tauT* (Fig. 2D). Taxonomic analysis revealed highest *tauE* expression in *Alphaproteobacteria* during the first bloom phase. An increase in expression was also observed during the second bloom phase, which was attributed to *Gammaproteobacteria* and *Flavobacteriia* (Fig. 2F). Further details about expressed genes for taurine utilization are provided in the Supplementary Information (Fig S2).

**Figure 2.**
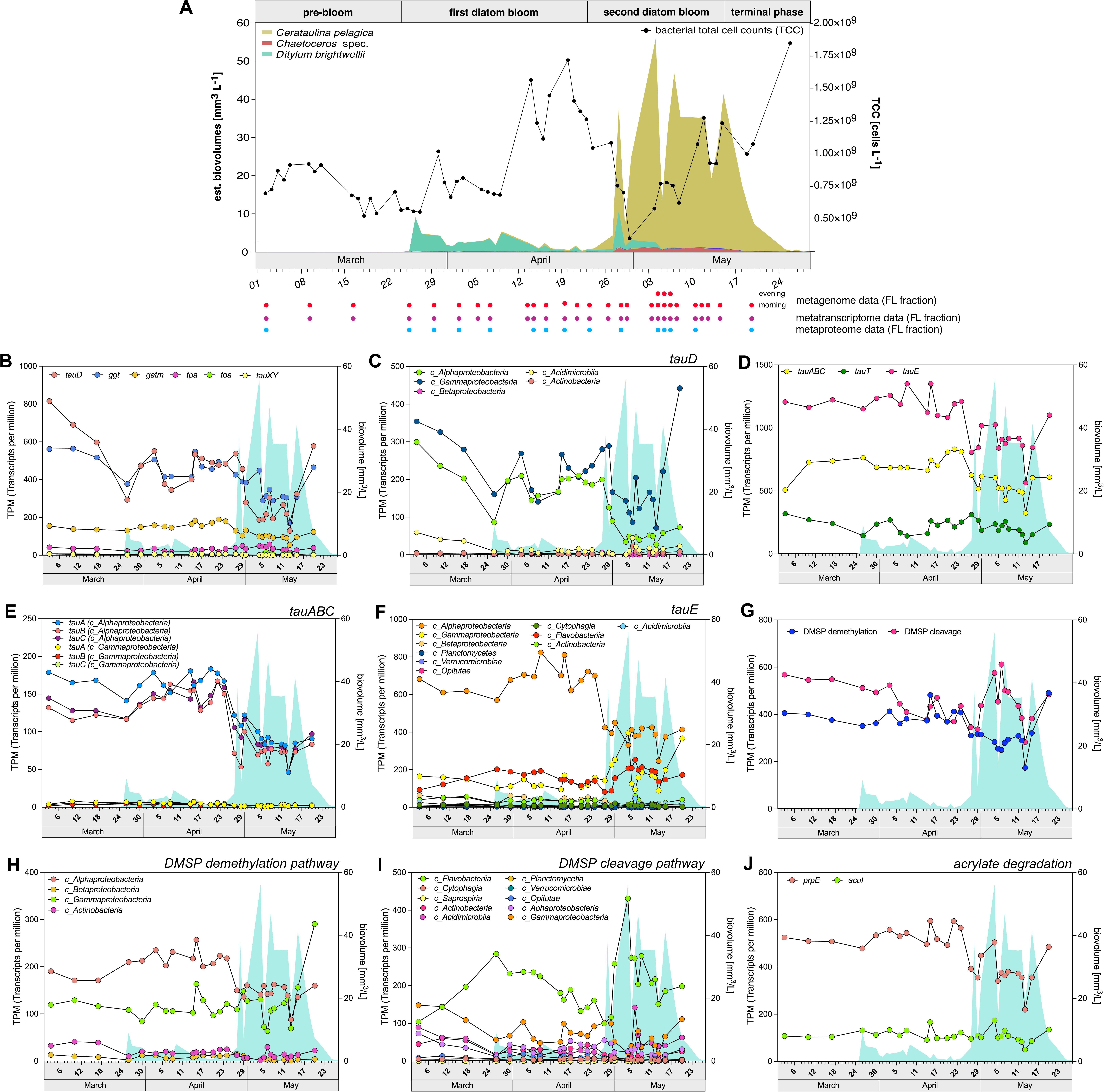
Expression analysis of genes involved in organosulfonate metabolism over the course of entire algal bloom of year 2020 based on assembled contig data. **A.** Schematic of the studied bloom (adapted from (13)). Only the three dominating diatoms are listed in the legend. Total bacterial cell counts were determined microscopically after staining with DAPI (4′,6-diamidino-2-phenylindole). Details have been published previously (13) **B.** Expression of genes involved in taurine degradation and transformation, **C.** taxonomic affiliation of taurine dioxygenase *tauD* gene; **D.** expression of taurine transporters; **E, F.** major clades attributed to taurine *tauABC* transporters and *tauE* expression; **G.** compiled expression of genes involved in DMSP demethylation and cleavage pathways; **H, I.** taxonomic affiliation of genes involved in both DMSP demethylation and cleavage pathways; **J.** expression of major genes *prpE* and *acuI* involved in marine acrylate cycle.

Regarding DMSP utilization, we found a drop in expression of DMSP demethylation pathway genes (*dmdABCD*) and an increase in expression of DMSP cleavage pathway genes (*dmsBC*, *ssuDE*, *tmoF*) during the second bloom phase (Fig. 2G). Taxa that expressed the former were mainly members of the *Alpha-* and *Gammaproteobacteria* classes, whereas the latter were expressed by members of the *Bacteroidota* phylum, mostly from the class *Flavobacteriia* (Fig. 2H,I). This suggest clear pathway preferences of these clades for DMSP utilization.

The DMSP cleavage product DMS is well-studied due to its presumed role in cloud formation, whereas less is known about the DMSP cleavage product acrylate and the marine acrylate cycle. Microbes capable of DMSP metabolism usually have genes to detoxify reactive acrylate (59). These genes include *acuI* (acrylyl-CoA reductase, K19745) and *prpE* (propionyl-CoA synthetase, K01908), the latter of which is also the initial enzyme in propionate utilization. We identified high overall *acuI* and *prpE* expression during both the first and second bloom phases (Fig. 2J). The expression of *prpE* during the first bloom phase could mainly be attributed to *Alpha-* and to a lesser extent *Gammaproteobacteria* and *Flavobacteriia*, however, during the second bloom phase, *prpE* expression declined in *Alphaproteobacteria*, while ramping up in *Flavobacteriia*, *Acidimicrobiia* (*Actinomycetota*) and *Gammaproteobacteria* (Fig. S1A-F). The more diagnostic *acuI* expression on the other hand was mostly attributed to *Alphaproteobacteria* with notable expression in *Gammaproteobacteria* and *Acidimicrobiia* during the second bloom phase (Fig. S1A-F), whereas no *acuI* gene was identified in *Flavobacteriia*. On overall, this demonstrates a widespread presence of these genes in various clades, indicating that acrylate is a readily available substrate during microalgal blooms. Moreover, the expression patterns of DMSP utilization pathways resembled those for acrylate conversion, consistent with the fact that acrylate results from the initial breakdown of DMSP.

Genes involved in the degradation of important C3-sulfonates include genes for the breakdown of dihydroxypropanesulfonate (DHPS) into sulfolactate (60) via hydrogenase enzymes (*hpsOPN*), the desulfonation of 3-sulfolactate by (2R)-sulfolactate sulfolyases (*suyAB*) and sulfoacetaldehyde acetyltransferase (*xsc*) (60, 61), and the decarboxylation of 3-sulfopyruvate to sulfoacetaldehyde (62) by sulfopyruvate decarboxylase (*comDE*). All these genes exhibited high expression during the first bloom phase (Fig. S1G,H), and were predominantly attributed to *Alphaproteobacteria* (Fig. S1). Further details about expressed genes for DMSP utilization are provided in the Supplementary Information (Fig S2).

### Seventy MAGs recruited about a third of all transcripts

As described previously (13), in total 251 species-level metagenome-assembled genomes (MAGs) of abundant FL bacterial clades were obtained from all metagenome data. Mapping of reads from corresponding 30 metatranscriptomes onto these MAGs recruited 41.4% of all transcripts, 80% of these transcripts were recruited by the 70 topmost expressed MAGs, corresponding to about a third of the total transcripts. Three of these MAGs were archaeal, devoid of organosulfur metabolism genes, and thus excluded from further analyses. Quality parameters, taxonomic affiliations and total expression values of the remaining 67 bacterial MAGs are summarized in Fig. 3.

**Figure 3.**
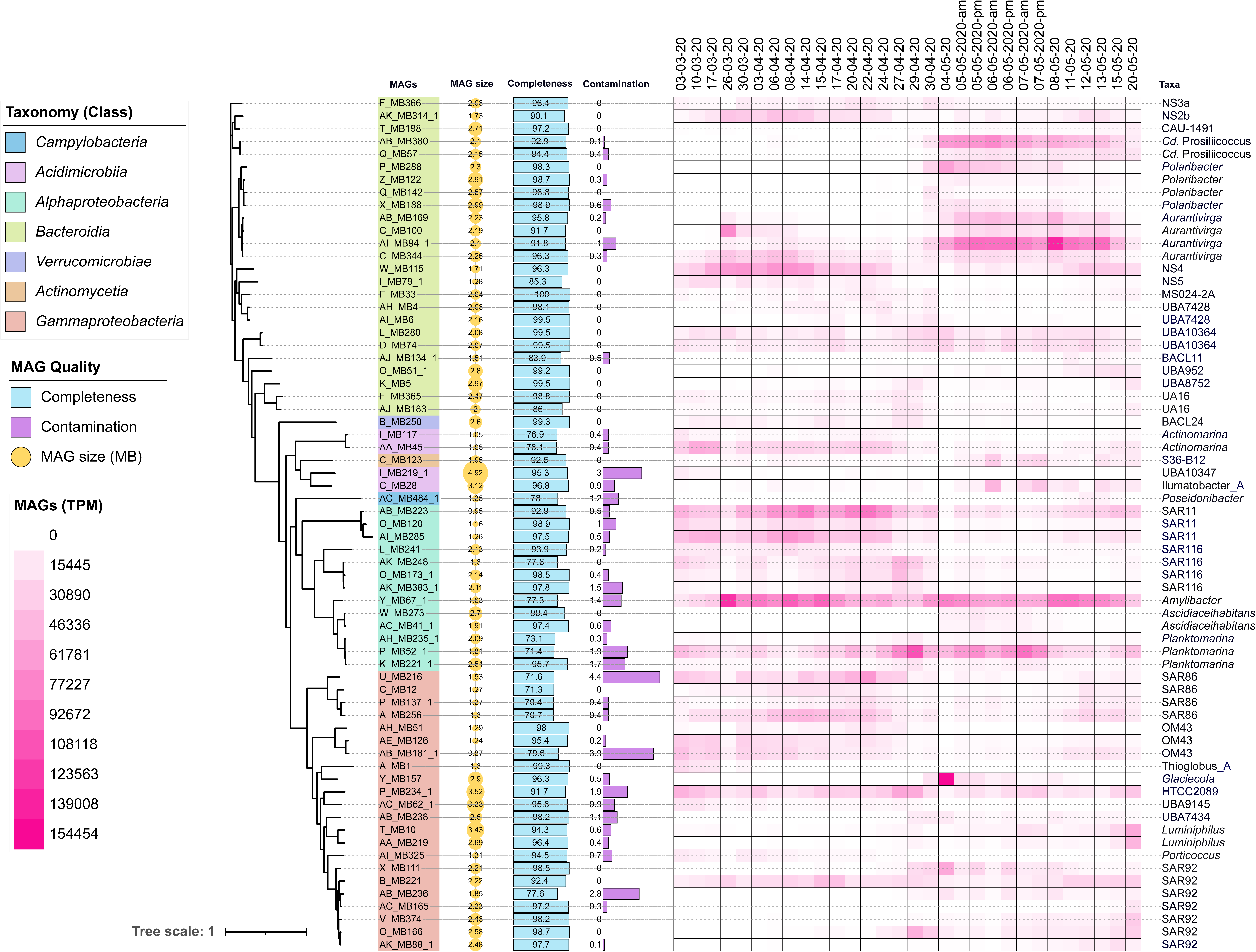
Topmost expressed 67 bacterial MAGs analyzed in this study. The phylogenomic tree was computed using *anvi-gen-phylogenomic-tree* within Anvi’o v7 (45). Completeness, contamination and taxonomic affiliations were calculated using checkM v1.0.18 (86) and GTDB-Tk v1.3.0 (87), respectively. The expression profile of all MAGs is represented as a heatmap of transcript per million (TPM) values along all 30 sampling points from March to May 2020. Tree annotations were done in iTOL v6 (88).

### Taurine was a substrate for many bacterial clades

Since our contig-based metagenome and -transcriptome analyses indicated high levels of taurine degradation, we analyzed the presence and expression of taurine utilization genes in the 67 topmost expressed bacterial MAGs. We found that taurine breakdown genes were wide-spread and present in 52/67 of the studied MAGs (corresponding KEGG categories are provided in Table S1 of the Supplementary Information). These 52 MAGs comprised genes for two different catabolic pathways: (i) the decarboxylating oxygenation of taurine to aminoacetaldehyde and sulfite via *tauD* (51, 52), and (ii) the oxidative transamination of taurine via *tpa*, *toa* and *xsc*. In addition, we also identified genes for the transformation of taurine to glutamyl-taurine by *ggt* (56) and to taurocyamine by *gatm* (57) (Fig. 4). We did not identify any *tauXY* taurine dehydrogenase genes in our MAGs, which some bacteria use to scavenge taurine-nitrogen (63).

**Figure 4.**
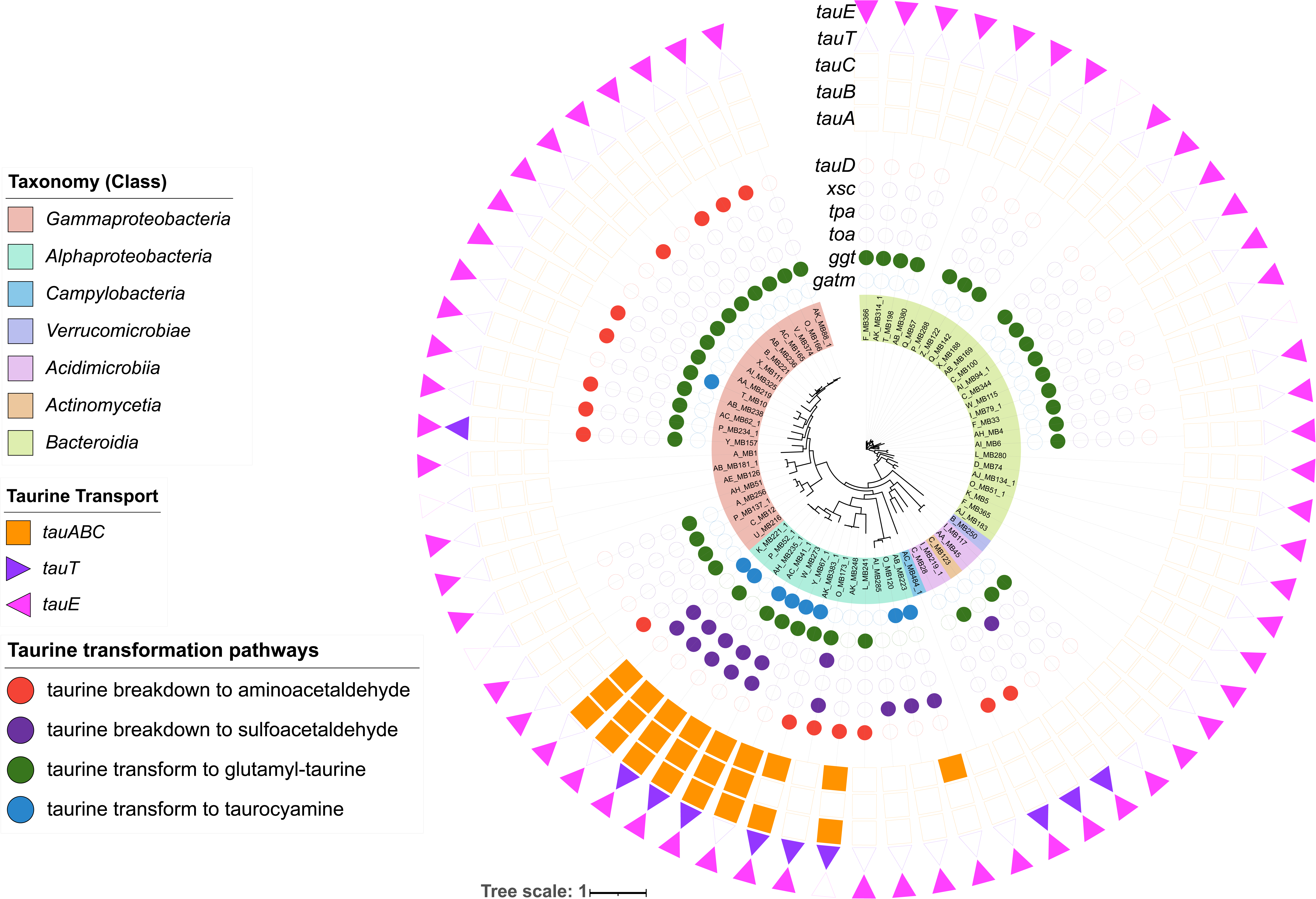
Presence of genes related to taurine breakdown/transformation and transport/uptake in the topmost expressed 67 bacterial MAGs in this study. The outmost ring denotes genus level taxonomic affiliations. Tree annotations were done in iTOL v6 (88).

In the 22 analyzed gammaproteobacterial MAGs, *tauD* was identified in ten, *ggt* in 18 and *gatm* in one MAG, including MAGs affiliating with *Luminiphilus* (T_MB10, AA_MB219), the SAR92 clade (AC_MB165, B_MB221, O_MB166, V_MB374), the SAR86 clade (U_MB216), *Glaciecola* (Y_MB157), the OM182 (P_MB234_1) and the UBA9145 (AC_MB62) clades. Furthermore, *tauD* was detected in acidimicrobial MAGs, specifically in MAGs belonging to the Ilumatobacter_A (C_MB28) and UBA10347 (I_MB219_1) clades. The latter harbored no less than 16 *tauD* instances, suggesting a particular relevance of taurine utilization. In contrast, within the examined alphaproteobacterial MAGs, *tauD* presence was restricted to MAGs affiliating with the SAR116 clade (Fig. 4), whereas members of the *Rhodobacteraceae* (except *Amylibacter*) possessed genes associated with the taurine transamination pathway (*tpa*, *xsc*). This underscores a clear predisposition of *Alphaproteobacteria* members to efficiently metabolize taurine via distinct metabolic routes. An exception was the SAR116 MAG O_MB173_1 (UBA3439 clade) with genes for both, taurine oxygenation (*tauD*) and transamination (*toa*, *xsc*). In stark contrast to *Alpha-* and *Gammaproteobacteria*, only *ggt* taurine transformation genes were found in MAGs of most of the abundant *Bacteroidota* (Fig. 4).

In terms of taurine-specific transport, *tauABC* genes were limited to *Alphaproteobacteria* and present in all six analyzed MAGs of *Rhodobacteraceae* members. In addition, *tauT* genes were present in actinomycetial (C_MB123) and acidimicrobiial (I_MB219_1 and C_MB28) MAGs (Fig. 4).

The metatranscriptome data revealed peaking *tauD* dioxygenase expression in SAR116 (MAG O_MB173_1) on April 27 (Fig. 5). Probably taurine sources were the blooming diatoms (64) and excretion products of zooplankters, such as copepods, whose abundances ramped up at the beginning of April and peaked around April 17 (weekly counting) (13). Presence of key proteins for taurine transport and utilization in *Alphaproteobacteria* could also be confirmed by metaproteome data, and included TauABC, Xsc, Tpa and Ggt, with highest protein abundance levels of Xsc in *Planktomarina* (MAG P_MB52_1), TauA in *Amylibacter* (MAG Y_MB67_1), SAR116 (MAG AK_MB248), and *Planktomarina* (MAG K_MB221) (Fig. S3).

**Figure 5.**
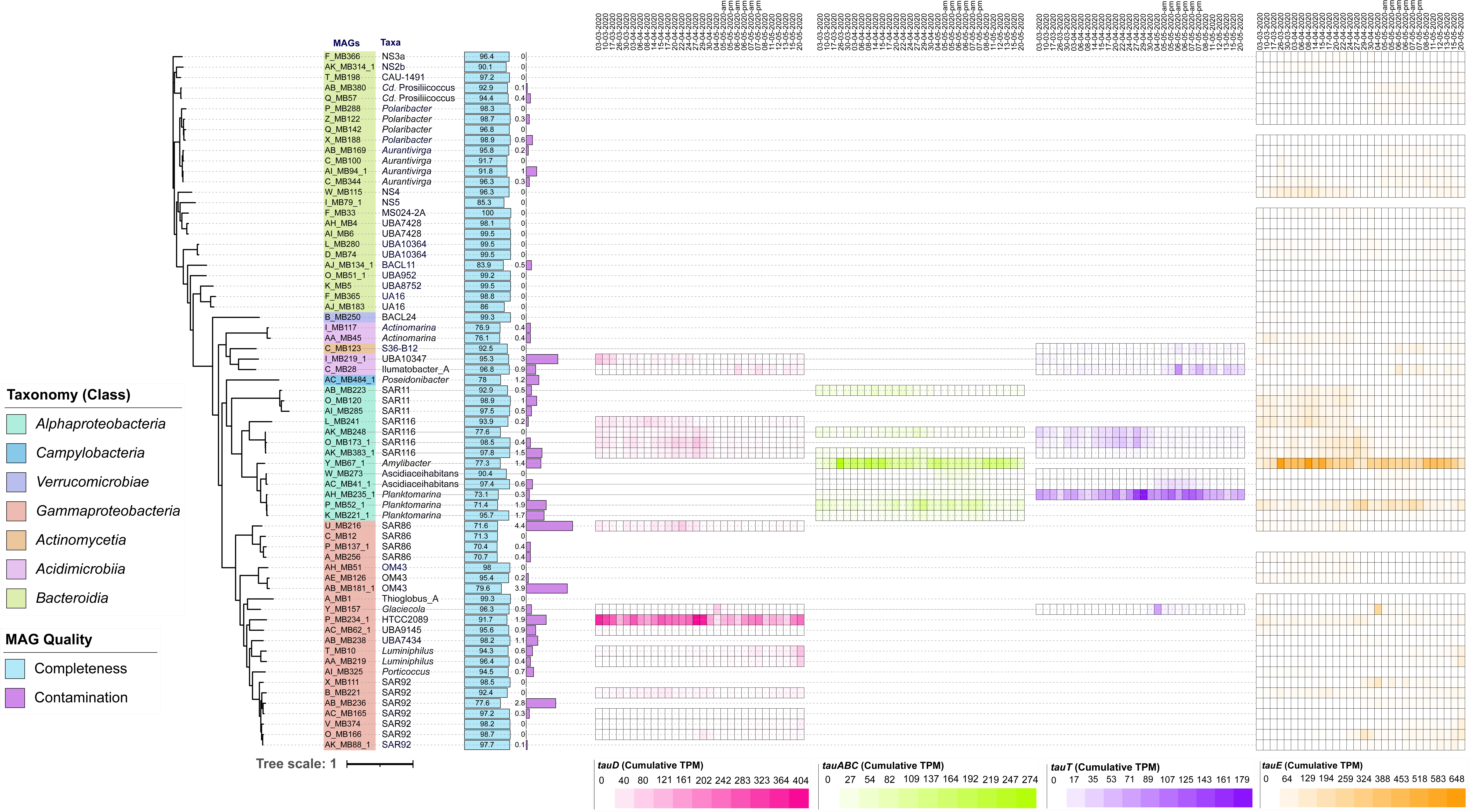
The expression profiles of genes related to taurine breakdown (*tauD* – shades of pink) and transport (*tauABC*, *tauT* and *tauE* – shades of green, purple and orange, respectively). Gene expression is depicted as heatmaps of TPM values along the course of the entire spring phytoplankton bloom of the year 2020 at Helgoland Roads. For MAGs with multiple instances of the same gene, the corresponding expressions were summated. Tree annotations were done in iTOL v6 (88).

Among *Gammaproteobacteria*, metatranscriptome data revealed the highest *tauD* expression in an OM182 clade member (MAG P_MB234_1) also with a peak on April 27 (Fig. 5). High expression was also observed in a *Glaciecola* member (MAG Y_MB157) with a peak on May 4, coinciding with the peak of the more intense second algal bloom phase. Interestingly, both *Luminiphilus* MAGs in the metatranscriptome dataset (T_MB10, AA_MB219) exhibited *tauD* expression during the terminal bloom phase, suggesting that these *Luminiphilus* members were scavenging taurine from diatom remains (Fig. 5). Metaproteome analysis also identified TauD in MAGs of SAR92 (O_MB166) and OM182 (P_MB234_1) representatives, whereas Ggt was confirmed in *Luminiphilus* (T_MB10, AA_MB219) and SAR92 clade (B_MB221) MAGs. Relative abundances of these protein groups in metaproteome data were considerably lower than those observed in *Alphaproteobacteria*.

### Methyl-sulfur metabolism genes were widely distributed among major bacterial classes

The 67 topmost expressed bacterial MAGs featured genes for both known DMSP utilization pathways (corresponding KEGG numbers and functional annotations are provided in Table S2 in the Supplementary Information). DMSP lyase genes were present in 62/67 MAGs, suggesting wide-spread DMSP utilization via cleavage among the most active bacterial clades during the bloom. Genes involved in the oxidation of the resulting DMS (*dmsBC*) and DSMO/DMSO_2_ (*ssuDE*) were identified in all 25 MAGs of *Bacteroidia* (*Flavobacteriia*). In the metaproteome data, we identified corresponding protein groups of 18 of the *Flavobacteriia* MAGs with high abundances in the sole NS4 clade MAG (W_MB115) during the first bloom phase (Fig. S3). During the more intense second bloom phase, metaproteome data showed high abundances of these protein groups in *Polaribacter* (X_MB188) and *Aurantivirga* (AB_MB169, C_MB100, C_MB344). Interestingly, we could not identify corresponding proteins for the on overall highest expressed *Polaribacter* (P_MB288) and *Aurantivirga* (AI_MB94_1) MAGs (13), but we did detect expression of the corresponding genes in the metatranscriptome data.

No DMSP demethylation pathway genes and proteins were identified in any of the *Flavobacteriia* MAGs (Fig. 6, S3), whereas MAGs of the gammaproteobacterial SAR92 and SAR86 clades exclusively featured genes and corresponding identified protein groups for the demethylation but none from cleavage pathways. These results corroborate our contig-based analysis and suggest distinct pathway preferences for DMSP utilization in the abundant *Flavobacteriia* and *Gammaproteobacteria* during the studied bloom.

**Figure 6.**
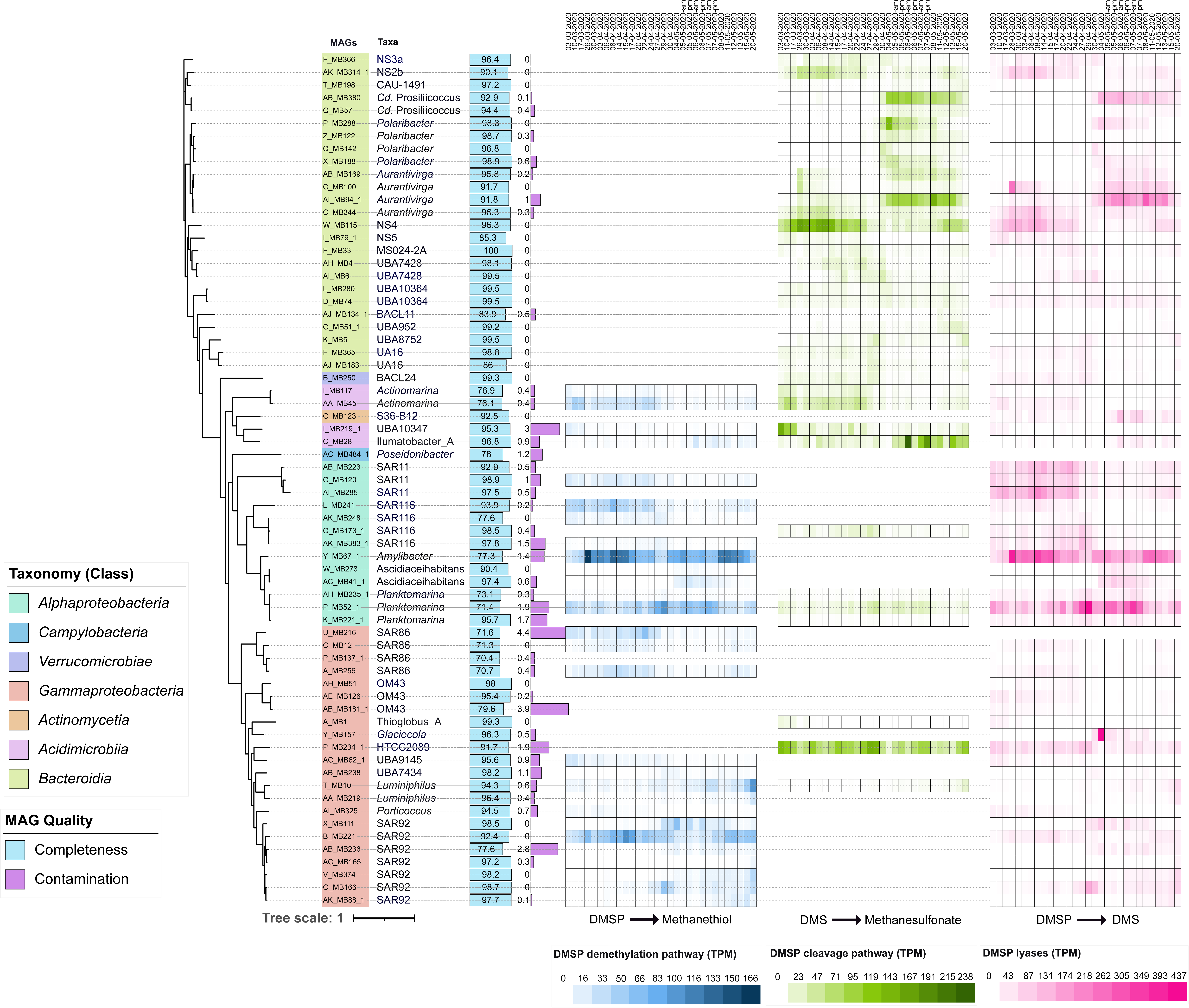
Expression profiles of genes involved in DMSP degradation, represented by heatmaps of TPM values. The genes for the demethylation pathway are depicted in shades of blue, whereas for those of the cleavage pathway are depicted in shades of green. DMSP lyases are plotted separately and are depicted in shades of pink. The KEGG and PFAM classes analyzed for these pathways are listed in the Supplementary Information. Tree annotations were done in iTOL v6 (88).

DMSP lyases transcripts were also identified in SAR86 (except U_MB216) and all seven SAR92 clade MAGs but only at low expression levels (Fig. 6). Conversely, only three of the gammaproteobacterial MAGs possessed genes for DMSP cleavage (P_MB234_1: HTCC2089 clade, A_MB1: *Thioglobus*_A, T_MB10: *Luminiphilus*), however, only HTCC2089 clade MAG P_MB234_1 exhibited expression in the metatranscriptome data. *Acidimicrobiia* MAGs (*Ilumatobacter*, *Actinomarina* and UBA10347) possessed genes of both cleavage and demethylation pathways, but expression levels of the former were about fourfold higher than those of the latter (Fig. 6). The UBA10347 clade MAG I_MB219_1 possessed fifteen instances of the *ssuD* and five of the *sfnG* gene, whereas *Ilumatobacter* MAG C_MB28 possessed four instances each, suggesting a high proficiency of these clades to utilize dissolved organic sulfonates. Genes for DMSP demethylation were identified in 28 of the 67 MAGs with highest expression in *Amylibacter* MAG (Y_MB67_1). A SAR116 member (MAG L_MB241) and *Luminiphilus* (MAG T_MB10) were found to contain all three *dmdABC* genes.

In terms of acrylate degradation genes, we identified *acuI* (acrylyl-CoA reductase, K19745) and *prpE* (propionyl-CoA synthetase, K01908) genes in the topmost expressed eight out of 13 alphaproteobacterial MAGs, two gammaproteobacterial MAGs (AA_MB219, AB_MB238) and both acidimicrobial MAGs (C_MB28, I_MB219_1). The expression of *acuI* was found to be highest in *Amylibacter* MAG Y_MB67_1 and coincided with the peak of the *D. brightwellii* microalgal population on March 26 (Fig. 7A). The expression of *prpE* was found to be highest on April 22 in SAR11 clade MAG AB_MB223, just before the onset of the second bloom phase (Fig. 7B).

**Figure 7.**
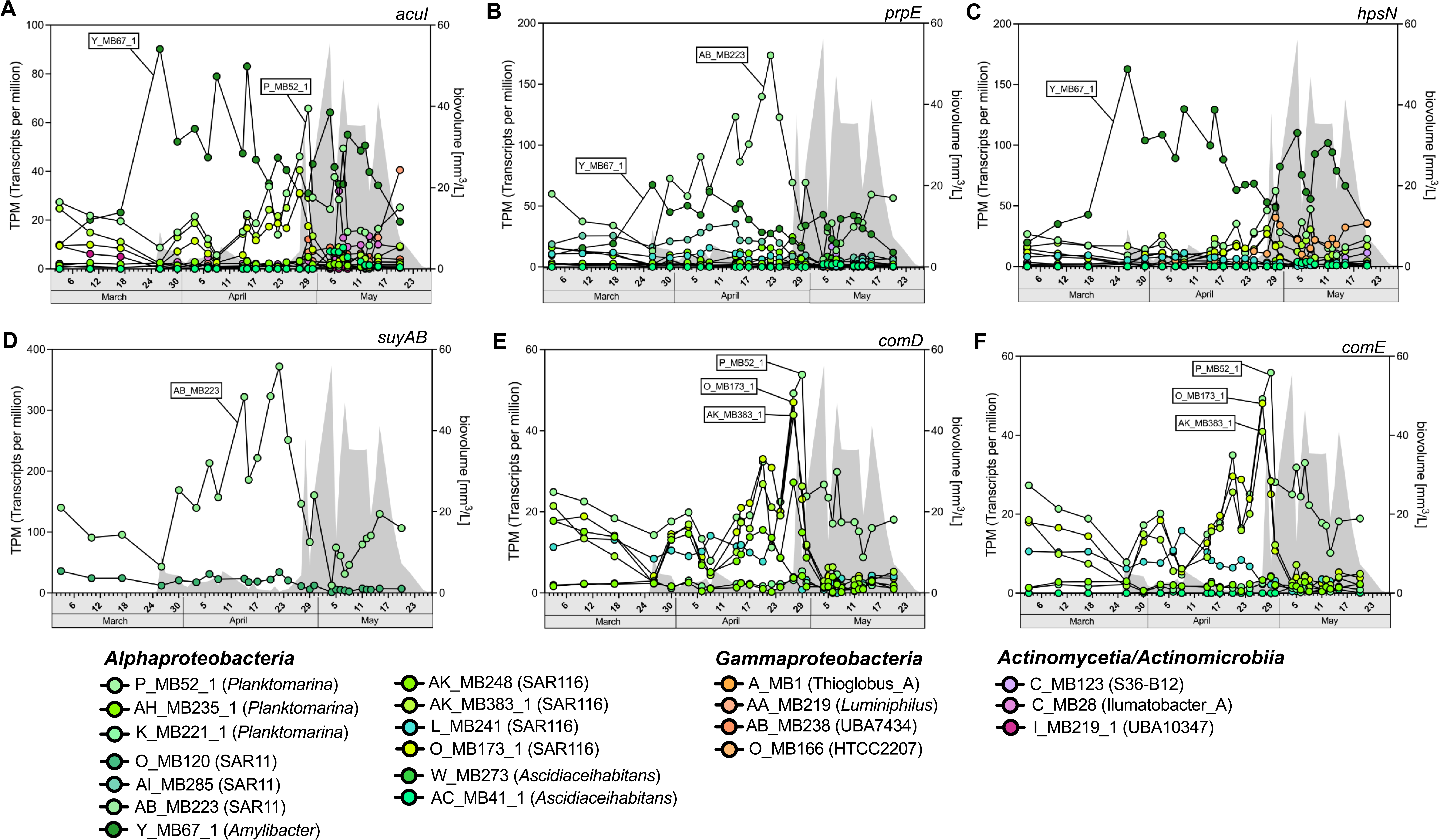
Expression profiles (represented by TPM values) of genes in top-expressed MAGs related to the utilization of methyl sulfur and other organosulfonate compounds over the course of algal bloom from March to May 2020 at Helgoland Roads. The estimated total algal biovolume (mm^3^/L) is plotted in the background (gray area). **A**. *acuI* (acrylyl-CoA reductase); **B**. *prpE* (propionyl-CoA synthetase); **C**. *hpsN* (sulfopropanediol 3-dehydrogenase); **D**. *suyAB* (sulfolactate sulfolyases); **E**. *comD;* **F**. *comE* (sulfopyruvate decarboxylase).

### Alphaproteobacteria utilized C3-sulfonates as a source of reduced sulfur

In aerobic bacteria, the degradation of the C3-sulfonate DHPS requires hydrogenase enzymes (*hpsOPN*) to produce sulfolactate (60) (Fig. 1). *HpsN* was found to be mostly restricted to alphaproteobacterial MAGs with occurrences in seven out of 13 alphaproteobacterial MAGs. The gene was highly expressed in *Amylibacter* MAG Y_MB67 with a peak on March 26 (Fig. 7C). This expression was followed by *Planktomarina* MAG P_MB52_1 and gammaproteobacterial HTCC2207 clade MAG O_MB166 in the later bloom stages.

The desulfonation of another C3-sulfonate, 3-sulfolactate, in bacteria is facilitated by (2R)-sulfolactate sulfolyases (*suyAB*) and *xsc*, resulting in the release of sulfur in the form of bisulfite (60, 61) (Fig. 1). One of three alphaproteobacterial SAR11 clade MAGs (AB_MB223) was distinct by showing high expression of genes involved in 3-sulfolactate desulfonation (Fig. 7D). Corresponding AB_MB223 SuyAB proteins were also identified in metaproteome data (Supplementary table S1). The other two SAR11 MAGs (AI_MB285, O_MB120) also featured mapped sulfolyase transcripts, but they were not among the highly expressed genes.

We also identified genes required for the decarboxylation of 3-sulfopyruvate to sulfoacetaldehyde by sulfopyruvate decarboxylase (*comD*, *comE*) (62) (Fig. 1). These genes were also restricted to all members of the alphaproteobacterial clades SAR116 and *Planktomarina*. The SAR116 MAGs (AK_MB383, O_MB173) each possessed three and two instances of the *comDE* genes, respectively. The highest expression of these genes was detected in *Planktomarina* MAG P_MB52, followed by SAR116 clade MAGs O_MB173 and AK_MB383 (Figs. 7E and F).

Detoxification of the sulfite resulting from enzymatic desulfonation reactions is crucial to avoid self-poisoning. One strategy involves direct sulfite export by dedicated transporters, such as TauE/SafE (K07090). Transcript analysis revealed expression of *tauE* in 62/67 MAGs (Fig. 4,5), with the MAGs of SAR116 clade members containing multiple instances. The SAR116 clade MAGs AK_MB383_1 and L_MB241 contained ten *tauE/safE* genes each, whereas MAG O_MB173_1 featured nine *tauE* genes. Notably, *tauE* genes were among the top 1% of expressed transcripts in all SAR116, *Amylibacter* (MAG Y_MB67_1) and *Planktomarina* (MAG P_MB52_1) MAGs.

## Discussion

Sulfur is essential to all life forms and present in various essential organic molecules, such as proteins, lipids, marine sulfated polysaccharides and enzyme cofactors. Phytoplankton has been estimated to annually assimilate about 1.36 Gt of sulfur into organic matter and to thereby support a major fraction of the oceanic surface sulfur fluxes (65). Phytoplankton-derived DOS compounds constitute an essential substrate for closely associated heterotrophic bacteria. In previous studies, we have demonstrated that phytoplankton-derived polysaccharides promote the proliferation of specialized microbial clades during spring algal blooms in the North Sea (12–14, 16, 47, 66). Here we use high temporal resolution deep metatranscriptome data in conjunction with state-of-the-art metaproteomics to show that also DOS compounds were utilized by abundant bacterial clades and thereby likely shaped the overall bacterioplankton compositional dynamics during the spring phytoplankton bloom at Helgoland in 2020. On overall, the expression of DOS transcripts aligned with MAG abundance profiles. For instance, the expression of the DMSP cleavage pathway, primarily attributed to members of the class *Flavobacteriia*, mirrored the abundance patterns of MAGs from the most abundant flavobacterial genera, *Polaribacter* (P_MB288) and *Aurantivirga* (AI_MB94_1).

Taurine is a common compound in mesozoo- and phytoplankton (45). Wide distribution of taurine utilization genes in highly expressed MAGs of FL bloom- associated bacteria supported the relevance of taurine during the studied phytoplankton bloom. For example, presence of 16 *tauD* genes in the acidimicrobial MAG I_MB219_1 (UBA10347 clade) suggest a high capacity for taurine utilization, potentially providing this clade with a competitive advantage over other taurine degraders. Such *tauD* genes were also identified in ten out of 22 gammaproteobacterial MAGs. On the contrary, not a single *tauD* gene was identified in any of the bacteroidotal MAGs. As we have shown previously, abundant and highly expressed members of *Bacteroidota* during the studied bloom specialized in the utilization of algal polysaccharides (13). This may explain the absence of taurine breakdown genes in the obtained largely complete bacteroidotal MAGs, as bloom-associated bacteria tend to exhibit pronounced resource partitioning as a strategy to thrive in a dedicated niche while avoiding competition. On overall, *Alphaproteobacteria* encoded the highest numbers of taurine utilization genes and also featured the highest diversity of taurine utilization pathways. For instance, the *tauD* gene was present only in SAR116 clade members, whereas members of the *Rhodobacteraceae* family and the SAR11 clade possessed transamination pathway genes (*toa*, *tpa*, *xsc*). While previous studies showed that SAR11 members can assimilate taurine (45), the exact biochemical routes and enzymatic processes in SAR11 members have not yet been elucidated. Our study suggests that members of the SAR11 metabolize taurine primarily through the transamination pathway rather than via taurine decarboxylation. Our results also revealed that taurine-specific transporters, such as TauABC and TauT, were exclusive to *Alphaproteobacteria*, underscoring the proficiency of *Alphaproteobacteria* for efficient taurine utilization. In contrast, although *Gammaproteobacteria* MAGs contained *tauD* genes, they lacked known taurine transporters, suggesting reliance on unspecific or as yet unknown mechanisms for taurine uptake. This likely reflects a less specialized more generalist approach of *Gammaproteobacteria* compared to *Alphaproteobacteria*. Besides phytoplankton, excretion products and fecal pellets of zooplankters such as copepods can be significant taurine sources (22, 41). This was likely the case during the studied bloom, where we observed a notable increase in copepod abundances in mid-April (for details, see (13)).

DMSP is the most well-studied marine sulfur metabolite and produced in large quantities by both micro- and macroalgae (30, 67, 68). It represents a substantial pool of dissolved organic sulfur in seawater. In marine phytoplankton, intracellular DMSP concentrations of up to 300 mM have been measured (69, 70). In contrast, the concentration of dissolved DMSP in open seawaters only occasionally reaches measurable levels, primarily due to its rapid remineralization by bacterial heterotrophs, particularly *Alphaproteobacteria* (71, 72). Presence of genes involved in the DMSP cleavage pathway has not yet been studied in *Bacteroidota*. Our results showed that all studied 25 bacteroidotal MAGs possessed genes for the oxidation of DMS and DMSO/DMSO_2_ along with DMSP lyases, with high expression during the second bloom-phase. This finding suggests that *Bacteroidota* represent another relevant group of DMSP consumers during algal blooms, expanding our understanding of their role in marine sulfur cycling. The ability of *Bacteroidota* to degrade DMS, DMSO, and DMSO_2_ may provide an energetic advantage by enabling them to derive extra energy, but could also serve as a source of reduced sulfur, especially when otherwise relying on marine polysaccharides that are poor in reduced sulfur. Gammaproteobacterial MAGs on the contrary did not possess genes for DMS/DMSO/DMSO_2_ oxidation but for demethylation. Likewise, most of the alphaproteobacterial MAGs also possessed demethylation pathway genes. By focusing on demethylation, these clades may effectively utilize DMSP as a resource without directly competing for the same sulfur intermediates. *Acidimicrobiia* MAGs possessed genes for both the cleavage and demethylation DMSP degradation pathways. However, the expression data indicated that only the cleavage pathway was active, suggesting a pronounced pathway preference in spite of the predicted genetic potential to employ both pathways. On general conclusion one might draw from our study is that *Flavobacteriia* and *Acidimicrobiia* members seem to preferentially utilize DMSP via the cleavage pathway, whereas *Alpha-* and *Gammaproteobacteria* are more inclined to utilize DMSP via DMSP the demethylation pathway.

Enzymatic cleavage of DMSP produces DMS and acrylate or 3- hydroxypropionate in equimolar concentrations (67). Another source of acrylate is the photolysis of DOM (73). Acrylate concentrations can reach up to micromolar levels in seawater, particularly during phytoplankton blooms (74). High expression of *acuI* in an *Amylibacter* MAG (Y_MB67_1) during the first bloom phase (March 26) suggests the diatom *D. brightwellii* as a probable source. Furthermore, the first bloom phase was characterized by top-down pressure from copepods, which may have led *D. brightwellii* cells to cleave their own DMSP in order to protect themselves via acrylate production, a grazing-activated defense mechanism that has been previously described for both bacteria (75) and microalgae (76). Besides, C3- sulfonates such as DHPS, 3-sulfolactate and 3-sulfopyruvate were found to be accumulated in high concentration inside phytoplankton cells (77).

On overall, our results indicate that members of the *Alphaproteobacteria* possess a high potential to degrade and utilize a variety of organosulfur metabolites during phytoplankton blooms, which confirms previous studies (71, 78, 79). We observed variations among species in the uptake and breakdown of specific sulfur metabolites, highlighting the diversity of organosulfur metabolism within *Alphaproteobacteria*. At the same time, *Gammaproteobacteria* and also some members of the *Bacteroidota* seem to utilize DOS compounds during phytoplankton blooms. Also, two of the studied acidimicrobial MAGs (C_MB28 and I_MB219_1) featured a remarkable potential to acquire reduced sulfur and carbon from organic sulfur compounds. Both, the roles of *Bacteroidota* and *Acidimicrobiia* in the metabolism of organosulfur compounds during phytoplankton blooms may have been underestimated and warrants an increased focus in future studies of phytoplankton bloom-associated bacteria and their role in sulfur and carbon cycling in marine systems.

### Study limitations

Contemporary long-read sequencing has enabled the retrieval of notable numbers of MAGs with medium to high quality from environmental samples of complex microbial communities. When coupled with metatranscriptomics and metaproteomics, these MAGs allow for as yet unprecedented insights into the metabolisms of abundant community members *in situ*. Still, even with high-quality long-read sequencing, MAGs remain bioinformatic constructs that can reflect consensus rather than actual genomes, in particular for species with high levels of strain heterogeneity. Fortunately, as we have shown in the past (80), fast-growing dominating species during phytoplankton blooms can be surprisingly clonal. However, we cannot exclude that some MAGs are not one-to-one reflections of actual genomes. In addition, while many of the MAGs in this study are of very high quality and largely complete, some are just of medium quality, which is why genes may have been missed. As valuable as the insights are from these types of environmental studies, they cannot substitute targeted isolation efforts of key players for detailed functional studies in reproducible laboratory experiments.

## Conclusions

In spite of the high innate complexity, we are step by step gaining a comprehensive understanding of the principles that govern bacterioplankton responses to phytoplankton bloom events. The various LMW organosulfur compounds that we explored in this study support bacterial heterotrophy and thereby shape the bacterioplankton community composition during such blooms. It is this highly evolved, adaptable yet specialized bacterial community that ensures that algal biomass is efficiently recycled. Hence, these bacteria are not only crucial for marine ecosystem functioning, but on a grander scale play an important role in the large carbon and sulfur biogeochemical cycles that regulate oceanic and atmospheric dynamics.

## Materials and Methods

### Sampling

Sampling, filtration and sequencing procedures were carried out as described previously (13). Briefly, samples were collected from March to May 2020 at the ’Kabeltonne’ long-term ecological research site (54° 11.3′ N, 7° 54.0′ E, DEIMS.iD: https://deims.org/1e96ef9b-0915-4661-849f-b3a72f5aa9b1). Seawater was sampled at 1 m depth on workdays for eleven consecutive weeks. Sequential filtration of ten liters of seawater was performed using 10 µm, 3 µm, and 0.2 µm pore-size 142 mm diameter polycarbonate filters. The filters were flash frozen in liquid nitrogen and stored at -80 °C until further use.

### Metagenomics, metatranscriptomics and metaproteomics

Metagenome and metatranscriptome sequencing of bacterial biomass from the 0.2- 3 µm size fractions was carried out at the Max Planck Genome Centre in Cologne, Germany, as described previously (13). A total of 30 metagenomes and corresponding metatranscriptomes were generated for selected time points based on chlorophyll *a* data and cell numbers (Fig. 2A). Metagenomes were sequenced on a PacBio Sequel II (Pacific Biosciences, Menlo Park, CA, USA) using one SMRT Cell per sample in HiFi long-read sequencing mode. Metatranscriptomes were sequenced on an Illumina HiSeq 3000 (Illumina Inc., San Diego, CA, USA) with about 100 million paired-end reads (2x 150 bp) per sample. The MAGs analyzed in this study were previously described in detail (13). In brief, all metagenomes were assembled individually using Flye v2.8.3 (81), binned and manually refined using Anvi’o v7 (82), which yielded 11,071 bins. These bins were dereplicated at 0.95 ANI (average nucleotide identity) to 251 representative MAGs, with quality parameters of min. 70% completeness and max. 5% contamination. Out of these 251 representative MAGs, 165 had > 90% completeness and < 5% contamination. Metaproteomes were analyzed on 15 dates for free-living samples (2020/03/10, 2020/03/26, 2020/03/30, 2020/04/03, 2020/04/08, 2020/04/15, 2020/04/17, 2020/04/20, 2020/04/24, 2020/04/29, 2020/05/05, 2020/05/06, 2020/05/07, 2020/05/11, 2020/05/20). Proteins were extracted from filtered biomass and subsequently analyzed (Supplementary Table S1). A detailed description of the metaproteome analysis is provided in the Supplementary Information.

### Omic data analysis

We employed an integrated analysis of metagenomes and metatranscriptomes using SqueezeMeta v1.3.1 (83) in merged mode as described previously (13). For, details on the generation of metagenome-assembled genomes (MAGs) and annotation, see (13). Mapping of processed mRNA reads to metagenome contigs was performed using Bowtie2 (84) and TPM values were calculated for each MAG as follows: (Σ reads of sample successfully mapping to a MAG x 10^6) / (Σ lengths of contigs of the MAG x Σ number of reads in the sample). Only the topmost 70 expressed MAGs were analyzed in detail for gene expression.

### Data availability

The metagenome, metatranscriptome, and MAG sequence data associated with this study can be accessed through the European Nucleotide Archive under the accession number PRJEB52999. Additionally, supporting environmental data can be found in the Zenodo repository, with the DOI 10.5281/zenodo.7656261. The mass spectrometry proteomic data have been deposited to the ProteomeXchange Consortium (http://proteomecentral.proteomexchange.org) via the PRIDE partner repository (85) with the dataset identifier PXD042805.

## Acknowledgements

We express our gratitude to Eva Maria Brodte, Antje Wichels, and Uwe Nettelmann from the Biological Station Helgoland (BAH-AWI, Germany), the long-term ecological research (LTER) site Helgoland team headed by Inga Kirstein, and the captains and crews of FS Aade and FS Uthörn for their invaluable assistance with sampling, analyses, logistics, and providing laboratory space. We furthermore express appreciation for Fengqing Wang, Mikkel Schultz-Johansen (MPI Bremen), Lilly Franzmeyer (University of Greifswald) for assisting sampling and Dr. Thomas Sura (University of Greifswald) for MS measurements.

This study was funded by the Max Planck Society and supported by the German Research Foundation (DFG) in the framework of the research unit FOR2406 ‘Proteogenomics of Marine Polysaccharide Utilization (POMPU)’ by grants of DB (BE 3869/4-2, BE 3869/4-3), TS (SCHW 595/10-3, SCHW 595/11-3), HT (TE 813/2-3), and RA (AM 73/9-3). The Helgoland Time Series of the Alfred Wegener Institute is supported by the Helmholtz Association as an LK-II performance category program.

## Footnotes

1 *Flavobacteriia* have recently been reclassified as *Bacteroidia*. Here we use the older taxonomy for clades within the *Bacteroidota* in order to ensure compatibility with previous publications.

